# Towards the Myoelectric Digital Twin: Ultra Fast and Realistic Modelling for Deep Learning

**DOI:** 10.1101/2021.06.07.447390

**Authors:** Kostiantyn Maksymenko, Alexander Kenneth Clarke, Irene Mendez Guerra, Samuel Deslauriers-Gauthier, Dario Farina

## Abstract

Muscle electrophysiology has emerged as a powerful tool to drive human machine interfaces, with many new recent applications outside the traditional clinical domains. It is currently a crucial component of control systems in robotics and virtual reality. However, more sophisticated, functional, and robust decoding algorithms are required to meet the fine control requirements of these new applications. Deep learning approaches have shown the highest potential in this regard. To be effective, deep learning requires a large amount of high-quality annotated data for training; the only option today is the use of experimental electromyography data. Yet the acquisition and labelling of training data is time-consuming and expensive. Moreover, the high-quality annotation of this data is often not possible because the ground truth labels are hidden. Data augmentation using simulations, a strategy applied in other deep learning applications, has never been attempted in electromyography due to the absence of computationally efficient and realistic models. Here, we present a new highly realistic and ultra-fast computational model tailored for the training of deep learning algorithms. For the first time, we are able to simulate arbitrary large datasets of realistic electromyography signals with high internal variability and leverage it to train deep learning algorithms. Because the computational model provides access to all the hidden parameters of the simulation, it also allows us to use some annotation strategies that are impossible with experimental data. We believe that this concept of Myoelectric Digital Twin allows new unprecedented approaches to muscular signals decoding and will accelerate the development of human-machine interfaces.

## 1 Introduction

Biosignals have been classically used for studying the underlying physiology, for clinical diagnostics, and for monitoring. More recently, they have also been used for interfacing humans with external devices. For example, signals measured at the surface of the skin from skeletal muscle electrical activity, i.e. surface electromyography (sEMG), are used for the control of bionic limbs [1]. In this application, the recorded electrical signals are converted into motion commands using machine learning [2, 3, 4]. In recent years, with the development of deep-learning based methods as well as wearable and cost-effective recording devices, there has been increased interest in using muscular signals as a basis for human-machine interfaces [5, 6]. The potential applications go well beyond the traditional clinical domains of prostheses and orthoses and range from robotic control to gaming and virtual reality [7]. A core challenge of deep-learning methods applied to biosignals is the acquisition of personalized and annotated training data in sufficient quantity and quality. Training data needs to be recorded for different subjects, at different times, with high variability in electrode configurations and experimental paradigms. In addition, it is challenging and in some cases impossible to properly describe the underlying physiological or neural parameters (e.g. individual muscle forces, fiber physiological parameters, motor neuron impulse timings), which are crucial for the correct annotation of data samples. As a result, acquiring experimental EMG data in sufficient quantity and quality is not only expensive and time-consuming, but in many cases not possible.

Data augmentation via simulation is an alternative approach to lengthy data acquisitions, and indeed augmentation techniques have been recently introduced for electrophysiological signals [8, 9, 10, 11]. However, most of these augmentation methods use “black-box” models, which aim to capture essential features of the signal without relating them to the underlying physiology [12]. Thus, the ground truth for most of the crucial parameters is still unknown, greatly limiting the potential use cases of such approaches. More sophisticated biophysical modelling methods are based on solving so-called forward equations (e.g., Poisson equation in the electrostatics case). However, this type of biophysical modelling has not been considered in the context of data augmentation for machine learning approaches. Indeed, state-of-the-art models are either not sufficiently realistic or not computationally efficient to produce suitable training data. For example, in the case of describing the generation of EMG signals, analytical models based on simple geometries of the tissues [13, 14, 15, 16, 17] provide simulations which reflect the broad characteristics of the signals, but cannot be used to reproduce specific experimental conditions due to the overly simplified anatomy. The more realistic models of EMG generation based on numerical solutions of the Poisson equation with generic volume conductor shapes [18, 19] are currently limited by their prohibitive computational time.

Here, we describe an EMG simulation method, based on the numerical solution of the forward equations suitable for deep learning data augmentation. It produces highly realistic EMG recordings, provides access to all underlying physiological parameters, and is extremely computationally efficient. Our results show that it is possible to simulate EMG signals for anatomically accurate conductor geometries and multiple muscles with tens of thousands of muscle fibers in a few seconds. As an application scenario, we also demonstrate the use of this model for data augmentation by pre-training neural networks that decompose EMG into the underlying neural activity sent from the spinal cord to muscles [20].

Our model is the only realistic and computationally efficient simulator targeted to AI training and approaching the concept of a Myoelectric Digital Twin. It allows generating arbitrary large datasets of realistic and personalized EMG signals, with high data variability and with a perfect annotation of diverse hidden parameters. As a result, our model may allow breakthrough approaches in AI-based EMG signal processing and decoding.

## 2 Results

### Biophysics

To allow the efficient simulation of a large quantity of highly realistic EMG recordings, we have developed a novel approach to solve the forward problem of the volume conductor in electrostatic conditions. Our approach is based on a hierarchical and flexible decomposition of the EMG simulation pipeline which allows the reuse and optimization of individual steps.

First, a realistic anatomy, described by bone, muscle, skin, and electrode surfaces, is discretized into a tetrahedral volume mesh. A conductivity tensor, anisotropic for muscles and isotropic elsewhere, is associated with each tetrahedral of the volume. Unlike the state-of-the-art approaches, which solve the quasi-static Maxwell’s equations for each fiber source and for each time instant, we solve them for a set of unit point sources located at each vertex of the mesh associated with the muscle tetrahedrals, which are referred to as basis sources. This computation does not depend on the time variable nor on the fibers and motor unit geometry and their physiological properties. *Therefore, changing these parameters does not require recomputing the forward solutions*.

Moreover, due to a rewriting of the equations involved using the so-called adjoint method, the solution is obtained by solving as many systems of equations as there are electrodes, rather than basis sources. Because the number of electrodes (≈ 10^2^) is typically much lower than the number of basis sources (≈ 10^5^), computational performance is substantially improved.

Second, using the same muscle surfaces used to describe the volume conductor, individual fiber geometries can be automatically generated, if this data is not available from other sources (e.g. from diffusion magnetic resonance imaging). Moreover, the fibers are grouped into motor units (MUs) following the state-of-the-art models for MU physiology. This step does not depend on the forward computations, and thus altering the related parameters and producing new simulation is highly efficient.

Third, the current source density propagating along the fibers is generated using a realistic intracellular action potential model. The contribution of individual fibers to the EMG recordings is obtained by discretizing each fiber into a set of points, integrating the current source density along its length, and projecting onto the sensor locations using the basis points computed in the first step. This approach effectively decouples the number of fibers and their discretization from the conductor model, allowing the simulator to handle tens of thousands of fibers per muscle. Again, changing the fiber parameters (end-plate location, action potential propagation velocity, tendons length, etc.) does not require recomputing the other blocks of the simulation.

Fourth, given a muscle activation profile, we use the size principle to recruit MUs and their associated fibers. This allows a simple and easily interpretable input to the simulation which can be used to simulate EMG recordings associated to specific muscle contractions and their movements.

The architecture described above, and detailed in Methods, has several advantages. First, each step of the procedure can be optimized individually, improving the performance of the system and the quality of the simulated EMG. In particular, due to the algebraic properties of the computations and their independence, a large part of them can potentially be performed in parallel (on CPU and GPU). Second, simulating data over a range of parameters does not require a full recomputation of the model. This allows the generation of massive EMG datasets covering a range of parameters and using personalized anatomy. In addition, the datasets are perfectly annotated, from overall muscle activation down to individual fiber action potential velocity.

As a result, our model is the first that allows the generation of ultra realistic and arbitrarily large (because of its computational performance) datasets of simulated EMG signals that can be used for AI training.

The details and all mathematical equations related to the model development are described in the Methods.

### The simulator reproduces analytical solutions

To produce realistic EMG data, the simulator leverages a flexible representation of the underlying anatomy and physiology. This flexibility does not only allow the use of realistic and personalized models, but also permits reproducing simple conductor geometry used in analytical solutions. A first validation of our numerical solution is performed by comparing it with its analytical counterpart for a cylindrical volume conductor geometry [21]. The normalized mean square error between the two solutions depended on the depth of the fiber and varied between 3% (1mm depth from the muscle surface) and 5% (11mm depth). Fig. 1 illustrates the analytical and numerical solutions for a fiber depth of 1 mm from the muscle surface. Because of the low error, the two waveforms are almost indistinguishable. It is important to note that the two volume conductor models in this validation are not identical. The theoretical/analytical solution is computed for an infinitely long cylinder (repeated periodically when discretized), while the numerical solution uses a cylinder of a large (sufficiently longer than the fiber and the electrode array), yet finite length. Increasing the length of the cylinder did not significantly alter the error.

**Figure 1:**
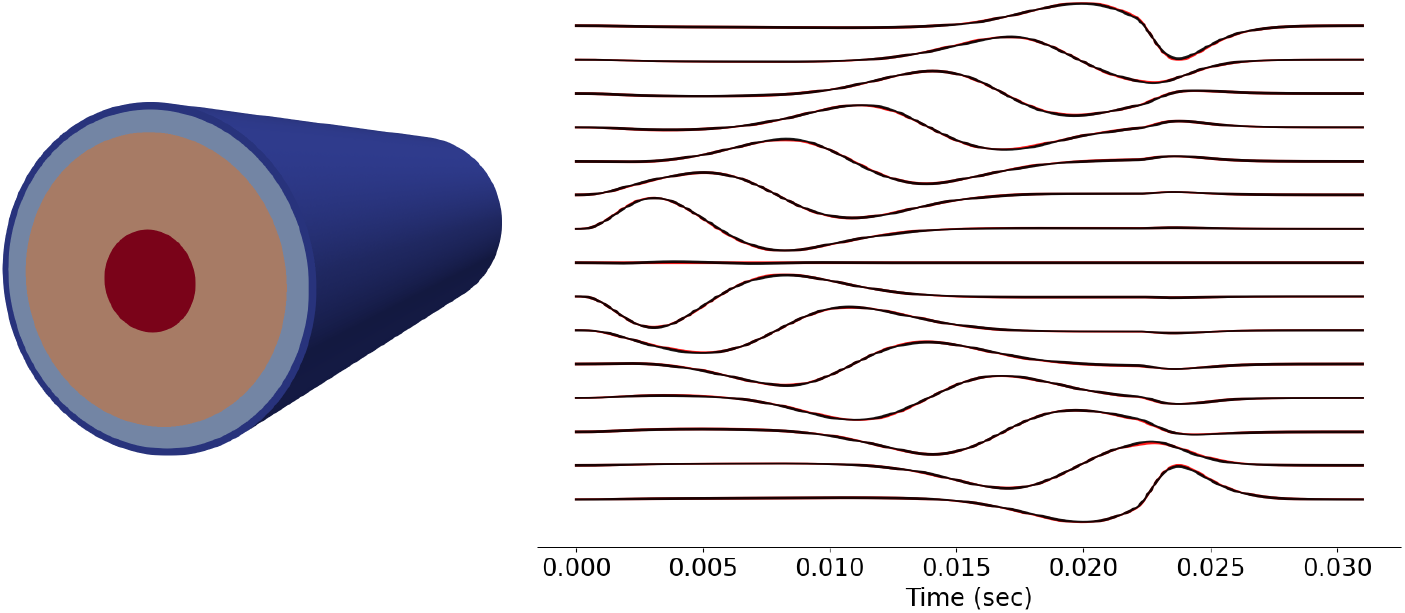
Comparison of the numerical and analytical [21] solutions (on the right) for a four layer cylindrical volume conductor model (on the left): analytical (red) and numerical (black) EMG signals for a differential array electrode montage. The depth of the source fiber in this example is 1 mm from the muscle surface.

### The simulator produces realistic EMG data

To evaluate the performance of the simulator at multiple scales, we started by simulating EMG signals associated to a single fiber activation inside the brachioradialis muscle. The signal recorded by an array of 16 rectangular electrodes (15 differential channels) when a single fiber was active is shown in Fig. 2A. The volume conductor model is based on an anatomically accurate forearm geometry, which includes all the muscles, bones, fat, and skin tissues.

**Figure 2:**
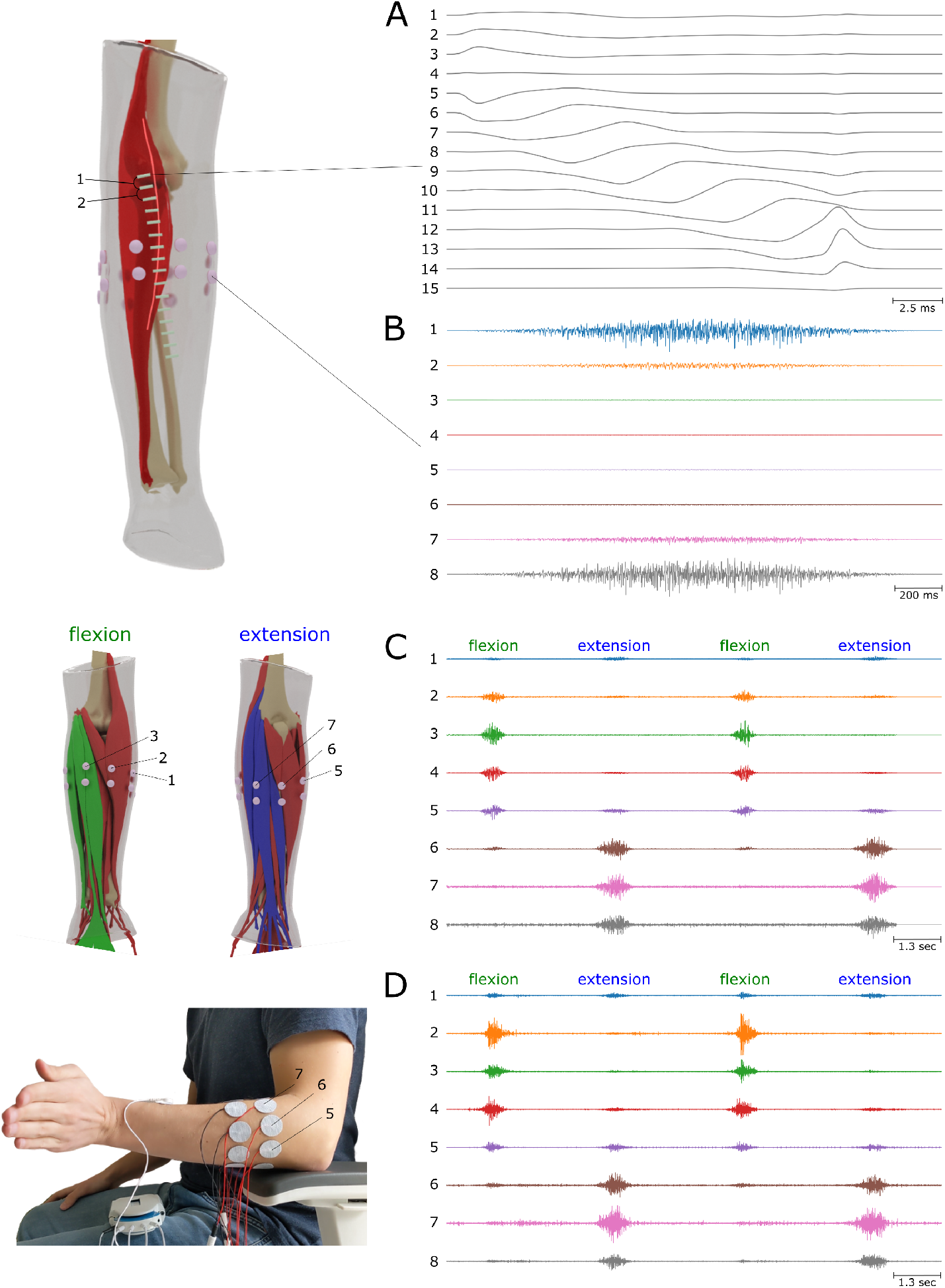
Simulation examples at multiple activation scales. (A) Single fiber activation in the Brachioradialis muscle measured by an electrode array with 15 differential channels. (B) 2-seconds long activation of the Brachioradialis muscle, reaching 100% of maximum voluntary contraction (MVC). 8 bipolar electrodes located around the forearm are simulated. (C) Simulation of wrist flexion and extension by activating the corresponding flexor and extensor muscles. (D) The experimental EMG signals of wrist flexion/extension.

Different distinctive features are present in the simulated signal that are also observed in experimental EMG signals [22]. In particular, electrodes of channel 4 are located on different sides of the neuromuscular junction (NMJ) and thus the respective signals cancel each other out. Channels 7-11 present propagating EMG components resulting from the fiber AP propagating from the NMJ to the tendons. Channels 2-6, as well as channels 12-15, contain non-propagating sEMG components, which are due to the AP generation at the NMJ and its extinction at the tendon (end-of-fiber effect), respectively.

A further example is a simulation of an excitation of a single muscle, illustrated in Fig. 2B. A simple excitation drive for the Brachioradialis muscle is simulated as gradually increasing from 0% to 100% of the maximum voluntary contraction and smoothly decreasing back to 0%. As described in Section 4.5, 50000 muscle fibers were realistically distributed into 200 motor units over the muscle volume and recruited according to the size principle [23]. The signal was simulated for 8 circular bipolar electrodes located around the forearm. In this example, the volume conductor effect becomes particularly visible with electrodes nearer to the active muscle having higher signal amplitudes. Notice that the electrodes record different signal waveforms as the muscle units are located at varying distances from the electrodes, weighting their contribution to the observed EMG signals. We also observe an increase of the signal amplitude with muscle excitation, an important feature of experimental EMG signals, which is a consequence of progressive motor unit recruitment and of an increase in the discharge rates of the active motor units.

Finally, we simulated sEMG signals from multiple muscle excitations, corresponding to the active wrist flexion and extension and passive wrist abduction against gravity. We used a simple muscle excitation model for three groups of muscles (flexors, extensors and abductors). More details about the experimental design are presented in Section Details of realistic simulation examples. Fig. 2C and Fig. 2D clearly show the qualitative similarities in signal characteristics between experimental and simulated data. Our model was able to reproduce the different signal patterns during both flexion and extension. Beside the different activation across the electrodes during flexion and extension, the effect of wrist abduction is also visible in both data sets. Thus, channels 2, 3 and 7, 8 present a small signal activity during the whole duration of the simulation, and not only during flexion/extension peaks. Similar activity can also be seen in experimental data, with channels 2 and 7 being the most active.

In addition to the analysis in the time domain, simulated data were compared against the experimental data in the frequency domain. Fig. 3 illustrates an example of the measured and simulated single channel sEMG. It has to be noted that the spectral characteristics of a signal strongly depends on multiple simulation parameters. In this example, we ran several hundreds simulations by varying the simulation parameters in a realistic range and selected the set of parameters leading to the minimal spectral difference. This approach, which is a simple version of inverse modelling, was possible because of the high computational speed of the simulations.

**Figure 3:**
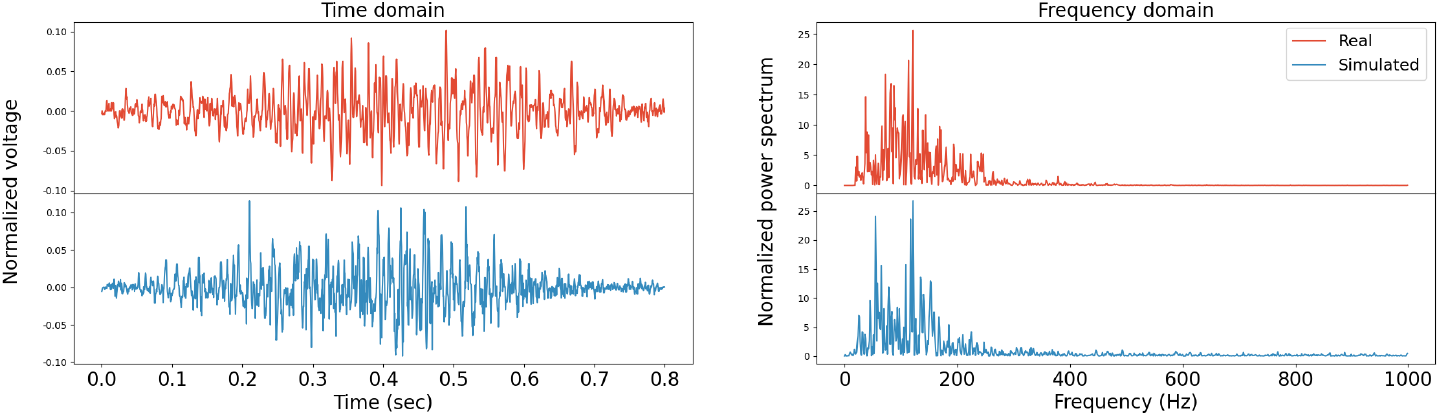
Comparison of experimental (red) and simulated (blue) single channel normalized sEMG signals in the time and frequency domains. The simulator parameters have been optimized to match the experimental signals.

### The simulator is ultra fast

The computational performance of an EMG signal simulation depends on the model properties and the particular experimental setup. Consequently, there is no benchmark to evaluate and compare the performance of different simulation methods. The computational time magnitude of the state-of-the-art methods is, in the best cases, in ***the order of hours*** for a single simulation (with a fixed set of model parameter values, ≈ 50000 fibers, 5 electrodes) [24, 19].

By exploiting the mathematical properties of the forward equations and source model, we were able to achieve a computational performance of ***the order of minutes*** per simulation. Moreover, in our model, changing most of the simulation parameters does not require recomputing the whole model and reduces the computational time of new simulations to ***the order of seconds***, if the volume conductor remains constant. As a result, it becomes practically possible to simulate arbitrary large datasets of highly realistic EMG signals with high variability in the simulation parameters. Details on the computational time in several conditions are provided in Methods (section Computational performance.

The proposed model is also highly scalable for multiprocessing, and the current computational time can be further reduced by several orders of magnitude by implementing parallel computation on CPU and GPU.

### Realistic and fast EMG simulations open unique perspectives for deep learning

Here, we show a potential use of high volumes of simulated surface EMG data for deep learning, utilising the proposed model to generate MUAP templates which can be used to pre-train neural networks. This methodology is used in other deep learning domains, such as the use of the ImageNet image database to pre-train object classifiers prior to adaptation to specific applications [25].

The myoelectric digital twin simulations (Fig. 4B) were used *to pre-train a neural network* that could extract motor unit activations from unprocessed HD-sEMG signal [26]. This pre-trained network was then trained to decompose experimentally measured HD-sEMG signals collected at the dominant wrist from nine participants (Fig. 4A). This procedure was then repeated, but with a randomly initialised version of the network instead of the pre-trained weights. See section 4.8 for details.

**Figure 4:**
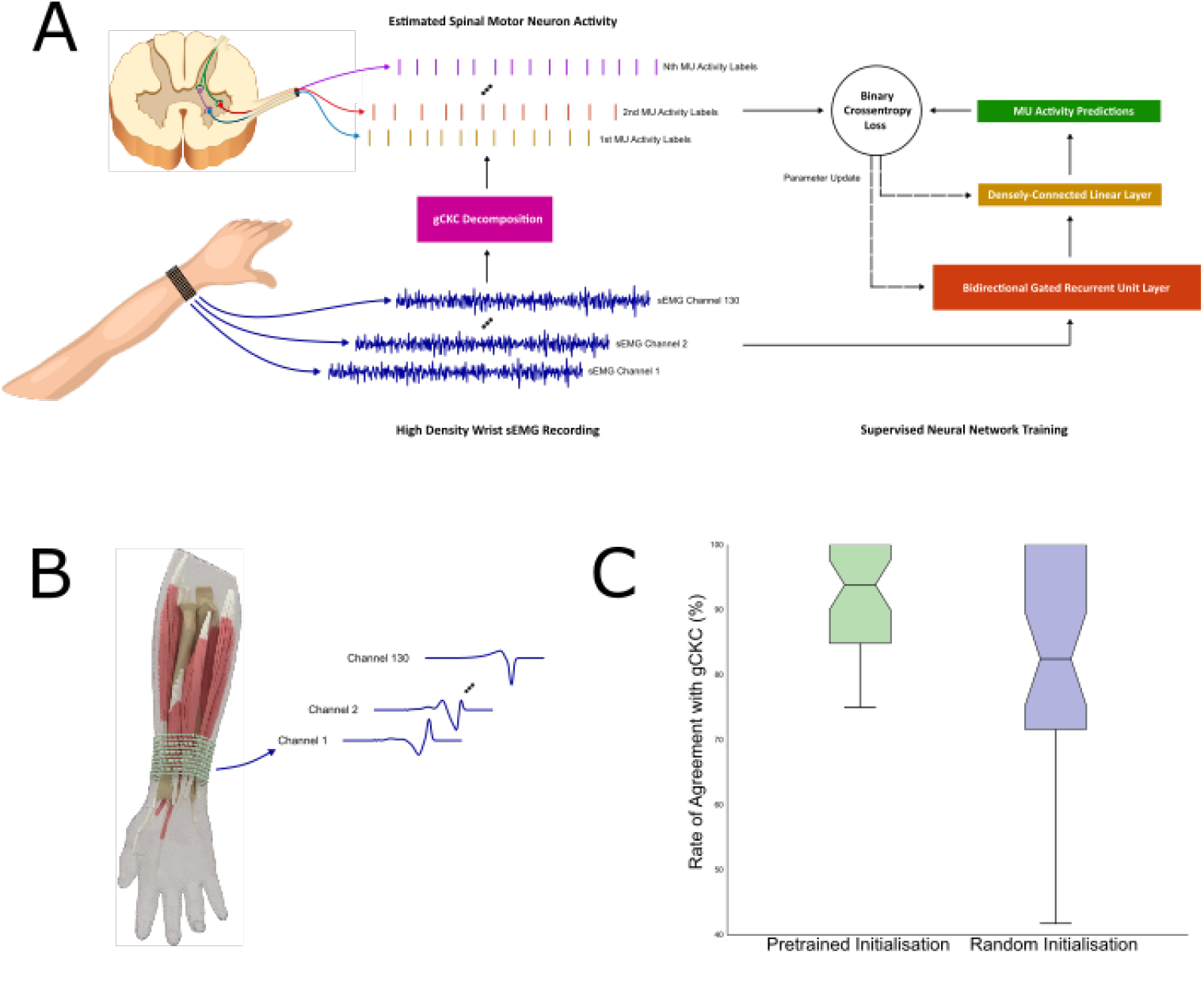
(A) Decomposition of experimental high density EMG recordings into underlying spinal motor neuron activities. The results obtained with the neural network (NN) were tested against the decomposition by a reference blind source separation method and manual editing by an expert operator. (B) Myoelectric digital twins were used to generate MUAP templates for different muscles and different model parameters (tissue conductivities, fiber properties, tendon sizes, etc). 64 sets, each containing 5 simulated MUAPs for 130 electrodes, were used for pre-training. (C) Rate of agreement (%) between the neural network MU activity predictions and the decomposition algorithm on one second of wrist flexor HD-sEMG signal. Median and interquartile range plotted over 39 motor units from nine participants. Both outputs were converted to timestamps using a two class K-means clustering. The neural network using a gated recurrent unit (GRU) network that was pre-trained using simulated EMG signal significantly outperformed a GRU with random initialisation (p *<*0.001).

The simulation pre-trained network outperformed random initialisation in decomposition accuracy when compared to the original decomposition as measured by the rate of agreement (RoA) metric [27] (Fig. 4C). The median (IQR) RoA of the pre-trained network was 93.8% (84.8 to 100.0), compared to 82.4% (71.6 to 100.0) in the random initialisation network, a significant difference according to the Wilcoxon signed-rank test (p *<*0.001). Of the 39 decoded motor units, 22 had improved RoAs with pre-training and one had a worse RoA, with the remaining 16 showing no change, generally because the initial RoA was already 100% without pre-training. The pre-trained network had a much lower variance in the accuracy of predictions on the test sets than random initialisation, quickly optimising to a model effective for generalisation to new signals.

## 3 Discussion

We have proposed an efficient computational approach to highly realistic surface EMG modeling. The method provides the solution to the generation of EMG signals from anatomically accurate volume conductor properties and number of muscle fibers, within limited computational time compatible with real-time signal generation. The proposed model is the only available EMG simulator with realistic description of the volume conductor and optimized for such computational efficiency. The main value of the model is that it eliminates bottlenecks of the state-of-the-art methods and opens unprecedented perspectives for using simulated sEMG for data augmentation in the deep learning framework and therefore for building a myoelectric digital twin.

The computational efficiency in the volume conductor solution has been recognized as an important component of EMG modeling, and some attempts to decrease the computational time in EMG simulations have been described. For example, the approaches developed by Dimitrov & Dimitrova [28] and Farina et al. [29, 21] substantially decreased the computational time in analytical EMG modeling for simple volume conductor geometries. These models provide simulations which reflect the broad characteristics of EMG signals, but can not be anatomically accurate because of the restrictions on the volume conductor and fiber source geometry. Realistic models using numerical solutions have also been recently proposed. The previous most complete and efficient model has been proposed by Pereira Botelho et al. [19]. These authors have used an anatomically accurate model to simulate EMG signals generated during index finger flexion and abduction. They gained computational speed by using the principle of reciprocity. In fact, one part of our calculations also includes the adjoint method, which is an algebraic representation of this principle. By using reciprocity, Pereira Botelho et al. [19] reported a computational time of 1 hour for simulating the activation of nearly 15500 fibers for 5 electrodes. This time, however, remains impractical for simulating arbitrary large data sets for a variety of parameter values. The model we proposed in this paper substantially surpasses the computational efficiency reported in [19]. We achieved it by efficiently exploiting mathematical properties of the forward equations, in particular by introducing the concept of basis points and by separating model parameters and variables into independent computational blocks. The approach does not only reduce the computational time for a full simulation, but also allows us to scale the solution, so that new solutions for the same volume conductor can be obtained without re-computing the volume conductor transformation. In this way, the generation of EMG signals within the same volume conductor, but varying all other simulation parameters, can be performed in extremely short time. Complex EMG signals from tens of thousands of muscle fibers located in multiple muscles, can be generated (and regenerated with different parameter values) in a computational time of the order of seconds. In contrast to previous models, our proposed simulator does not compromise accuracy and computational speed.

Some limitations remain in the current state of the presented model. It does not include some sources of variability that are present in experimental EMG signals and strongly impact their processing and analysis. For example, the model does not include advanced noise and artifacts descriptions, biomechanical modeling of the musculoskeletal system, and non-stationary volume conductor properties and fiber geometry. While these aspects are beyond the scope of this paper, they are relevant features to include in future developments.

The advances presented in this work, together with the proposed future developments, naturally lead to the concept of a myoelectric digital twin - a hyperrealistic, personalized, computationally-efficient model which generates EMG data in a quality and quantity sufficient not only to augment but to replace real data, with utility for AI training in the various real world applications. Here we have illustrated the potential of this approach by augmenting training data for deep neural networks, with the aim of identifying the discharge times of spinal motor neurons from surface EMG signal. By using the simulator to augment training (through a pre-training procedure), we showed a substantial increase in the performance of the decomposition network when applied to experimental data, demonstrating a highly relevant use of the proposed approach for decreasing the need for experimental training data in human-machine interfacing applications.

## 4 Methods

### 4.1 Forward problem

The fiber extracellular potentials that are measured by EMG electrodes are generated by transmembrane currents. The properties of bioelectric currents and potential fields can be determined from solutions of the Maxwell’s equations, taking into account the electrical properties of biological tissues. Because of the relatively low frequencies of signal sources of biological origin, the quasi-static assumption can be applied [30, 31], so that the electric potential and the primary current sources are related by the following Poisson equation [30, 32, 33] with Neumann boundary conditions:

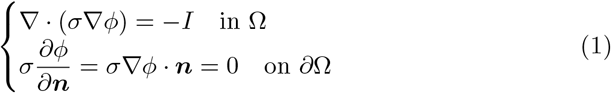

where Ω ⊂ ℝ^3^ is a volume conductor domain of interest, *∂*Ω its boundary with outward pointing normal unit vector ***n***, *φ*(***r***) [*V*] is the electric potential, *I*(***r***) [*A/m*^3^] is the current source density (CSD), *σ*(***r***) [*S/m*] is a conductivity tensor. The second line of the equation (boundary condition) reflects the assumption that no current flows out of the domain of interest. In the context of EMG modeling, this implies that there is no current flow between the skin and air. The current source density *I*(***r***) is interpreted as the volume density of current entering or leaving the extracellular medium at position ***r*** ∈ Ω. A negative CSD corresponds to current leaving the extracellular medium (due to the fiber transmembrane currents) and is thus conventionally called a sink. Likewise, current entering the extracellular medium is called a source [34, 35].

Equation (1) cannot be solved analytically for general volume conductor geometries, but several numerical methods can be used to approximate its solution. Here, we use the finite element method (FEM) [36], which discretizes the volume conductor Ω as a tetrahedral mesh Ω_*t*_. Given this mesh, we use the Galerkin method to project the potential *φ* onto the space of piecewise affine functions defined on Ω_*t*_. Fig. 5A and Fig. 5B illustrate an example of a realistic forearm model and corresponding discretized volume mesh respectively.

**Figure 5:**
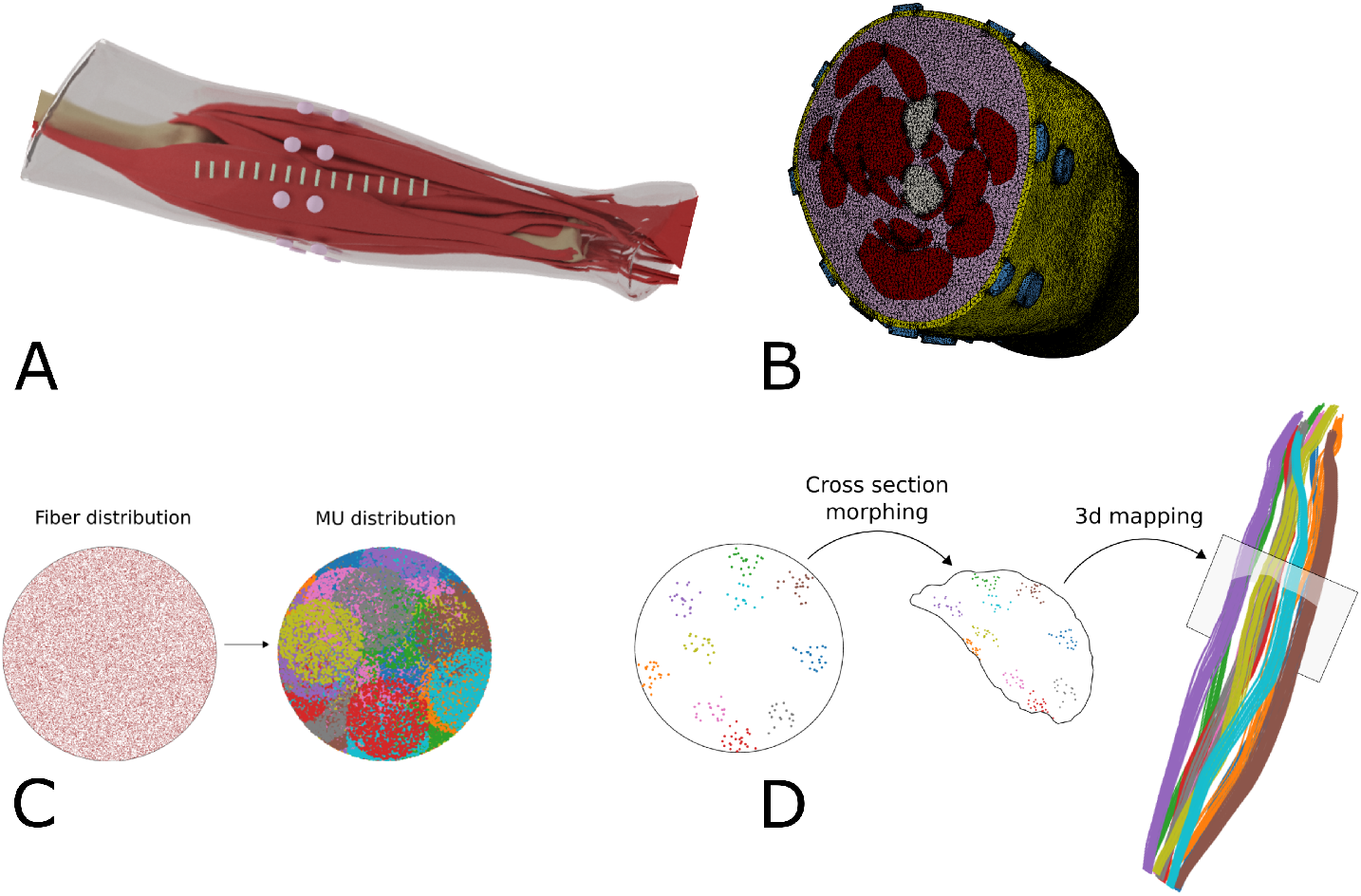
(A) Surface geometry of muscles, bones, subcutaneous tissue, skin and electrodes used for arm modeling (taken from BodyParts3D, The Database Center for Life Science (http://lifesciencedb.jp/bp3d/)). (B) Cross-section of the volume mesh generated from the arm surfaces. (C) Uniformly distributed fibers inside a unit circle are grouped into motor units of different sizes, locations and territories. (D) Example of mapping of 10 small motor units from the circle into an arbitrary muscle by morphing the unit circle into the muscle cross section.

This discretization process converts the continuous operator problem of Eq. (1) to a finite system of linear equations:

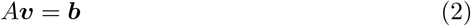

where *A* is a symmetric and sparse *n*_*v*_ × *n*_*v*_ matrix, *n*_*v*_ is the number of mesh vertices, 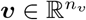 is a vector of potential values at mesh nodes, and 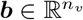 is a vector containing source information. Because the electric potential is defined up to a constant, the matrix *A* always has a one dimensional null space. To obtain a unique solution to the system of Eq. (2), we constrain potentials ***v*** to have a zero sum.

In the context of EMG, we are not interested in finding electric potentials everywhere in the conductor, but only at the electrode locations. Let *S* be a selection matrix with a shape *n*_*e*_ × *n*_*v*_ which only selects the values at EMG electrode locations (*n*_*e*_ is the number of electrodes). Each row of *S* can be designed to select a single point location or to integrate over an area (e.g. the electrode-skin interface) depending on the location and number of its non-zero elements. Also, let ***b***(***r***) correspond to a point source at location ***r***. The resulting EMG signal is thus given by:

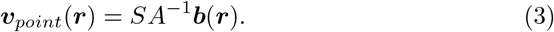

Let us analyze in more detail the structure of *A* and ***b*** from Eq. (2). Let {*w*^*i*^(***r***), *i* = 1…*n*_*v*_} be a set of *n*_*v*_ *P* ^1^ (piecewise linear) basis functions over the tetrahedral mesh Ω_*t*_. Note, that *w*^*i*^ is 1 at the *i*-th vertex of the mesh, is 0 at all other vertices and is linear at all tetrahedra adjacent to the *i*-th vertex. In this case, *A* and ***b*** have the following structure:

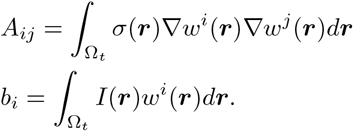

First, let us notice that *A* is symmetric and, in general, a very large matrix which can be stored only because it is sparse. Indeed, the functions *w*^*i*^ have a compact support and their pairwise scalar product is non-zero only for “neighbor” functions. Since the pseudo-inverse (or the inverse) of a sparse matrix is usually not a sparse matrix, it is impractical to compute it because of the amount of memory needed to store it. Thus, iterative methods are typically used to solve the system of Eq. (2) for every given ***b***.

Consider the case of 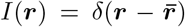 which corresponds to a unit point current source at location 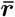. Without loss of generality, we assume that this source is inside a tetrahedron formed by the vertices *i*_1_, …, *i*_4_ of the mesh. In this case, we obtain:

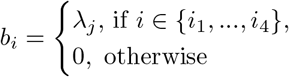

where {*λ*_*j*_, *j* = 1, …, 4} are the barycentric coordinates of the point 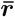 inside the tetrahedron {*i*_1_, …, *i*_4_}. Applying this expression to Eq. (3), we obtain:

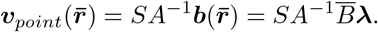

where 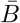 is a *n*_*v*_ × 4 matrix with 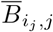 for *j* = 1, …, 4, and 0 otherwise. This implies that the solution of the system of Eq. (2) for any unit point source can be computed as a barycentric sum of solutions on the vertices of the corresponding tetrahedron. Therefore, it is sufficient to compute solutions of Eq. (2) for “basis” sources located on mesh vertices, to be able to evaluate a solution for any point inside this mesh efficiently. Let *n*_*s*_ be the number of such basis sources. For the most general case, when the source can be located anywhere inside the mesh and *n*_*s*_ = *n*_*v*_, let *B* be a *n*_*v*_ × *n*_*s*_ identity matrix. The objective is to compute “basis” solutions:

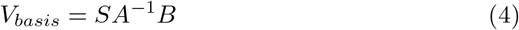

where *V*_*basis*_ is a *n*_*e*_ × *n*_*s*_ matrix, whose columns contain the solutions of Eq. (2) for a unit point source located at the corresponding mesh vertex. Hence, the potentials for any source location ***r*** is given by:

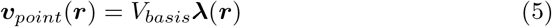

where 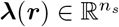 is a vector, whose four non-zero elements contain the barycentric coordinates of point ***r*** inside a corresponding tetrahedron. Note, that one may restrict potential sources to be located inside specific subdomains of the whole mesh (which is the case for EMG). In this case, *n*_*s*_ corresponds to the number of vertices of these subdomains, and the matrix *B* is a submatrix of the identity matrix.

The most straightforward way to compute *V*_*basis*_ from Eq. (4) is to solve a problem of the form *A****x*** = ***b***_***i***_ for each column of the matrix *B*. It would thus require solving *n*_*s*_ systems of linear equations. For realistic conductor geometries, which have a large number of vertices, solving a single system may take up to a few minutes and solving *n*_*s*_ systems quickly becomes impractical. Therefore, we propose the use of the adjoint method [37], which requires solving *n*_*e*_ systems only. In the context of EMG, the number of electrodes is usually significantly smaller than the number of vertices in the muscle subdomain meshes, i.e. *n*_*e*_ *<< n*_*s*_. Let us define *K* = *SA*^−1^, which is a matrix of size *n*_*e*_ × *n*_*v*_. Because *A* is symmetric, and the inverse of a symmetric matrix is also symmetric, we can write *K*^*T*^ = *A*^*−*1^*S*^*T*^. Then, *K* can be found by solving the system:

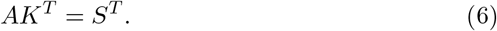

The matrix *S*^*T*^ has *n*_*e*_ columns and, thus, only *n*_*e*_ linear systems need to be solved to find *K*. The basis solutions can then be found as:

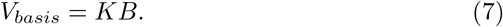

### 4.2 EMG signal of a single fiber activation

The action potential generated by the flow of ionic currents across the muscle fiber membrane is the source of excitation. For a given intracellular action potential (IAP) model *V*_*m*_(*z*), the transmembrane current source per unit length is proportional to the second derivative of *V*_*m*_(*z*), where *z* is a fiber arc length measured in *mm*. A general description of the current source density traveling at velocity *v* along the fiber with the origin at the neuromuscular junction at location *z*_0_ is [27, 29, 38]:

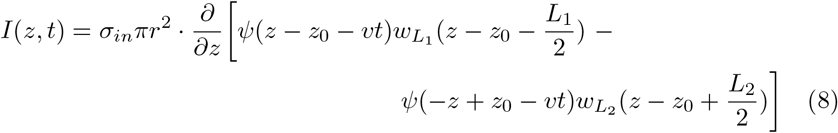

where *z* ∈ [0, *L*] is a location along the fiber of length 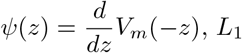, *L*_1_ and *L*_2_ are the semi-lengths of the fiber from the end-plate to the right and to *mm* the left tendon, respectively, *σ*_*in*_ is the intracellular conductivity, and *r* is the fiber radius. We have chosen *w*_*L*_ to be a Tukey window, as proposed in [24]. The IAP 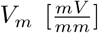 can be mathematically described in the space domain as proposed in [39]:

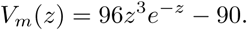

Let ***r***(*z*) be a fiber geometry parametrized with respect to the fiber arc length *z*. Combining the transfer function of a point source in Eq. (3) with the fiber’s current density in Eq. (8), we obtain the equation for the EMG signal resulting from a single fiber activation:

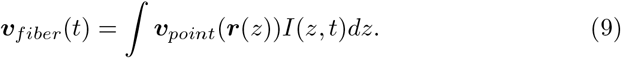

This integral can be efficiently approximated by discretizing the fiber geometry into sufficiently dense spatial samples {***r***(*z*_*i*_)}_*i*_ and assuming that ***v***_*point*_(***r***(*z*)) is piecewise constant around these points. If we also rewrite Eq. (8) in a shorter form as 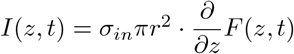, Eq (9) becomes

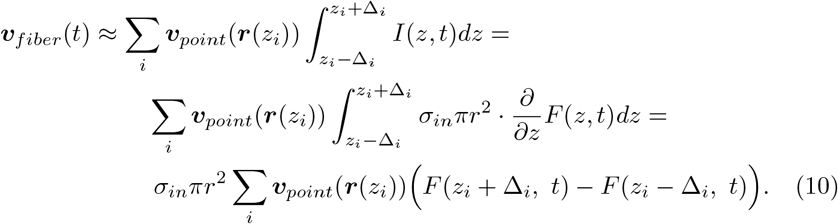

Note, that ***v***_*point*_(***r***(*z*_*i*_)) can be efficiently computed from Eq. (5). Moreover, once ***v***_*point*_(***r***(*z*_*i*_)) are computed for all given fibers, we can change the parameters of the current source density (action potential waveform shape, propagation velocity, location of neuromuscular junction), and compute the corresponding EMG signal with Eq. (10) by only matrix multiplication complexity.

### 4.3 Geometrical and physiological modeling of motor units

The motor unit action potential (MUAP) is the summation of the single fiber action potentials (APs) of the muscle fibers in the MU. Different types of MUs can be modeled [40, 41]. Our approach consists in generating fiber and motor unit distributions inside a unit circle, and then projecting it into arbitrary 3D muscle geometry (Fig. 5D), using methods similar to those described in [42]. This provides a high level of control for the fiber and MU distribution parameters independently of a particular muscle geometry. A common way to simulate fibers and MUs is to start by defining MU positions, sizes and territories, and then simulate fibers inside these MUs according to their parameters [43, 44]. We, however, propose another approach. First, we simulate uniformly distributed fibers inside a unit circle. Then, MU centers and their circular territories are generated and, finally, we associate each fiber to an MU. A fiber is associated to one of the MUs that contains it inside its territory with a probability proportional to the MU density (Fig. 5C). This approach has two main advantages. First, it guaranties (by construction) the uniform fiber distribution inside a circular muscle cross-section. Second, once fibers are generated and projected into a muscle geometry, different MU distributions can be generated very quickly, without regenerating fibers and recomputing transfer functions ***v***_*point*_(***r***(*z*_*i*_)) for their nodes.

#### MU recruitment model

During muscle contraction, the MUs are recruited according to the size principle [23]. This can be simulated by associating a threshold of excitation to each MU, as described for example by Fuglevand et al. [45]. Linear or non-linear rate coding models can be used [45, 46, 47].

The excitation rate as a function of time for each muscle is converted into the firing rates of the active MUs. Inter-discharge intervals are then generated with variability of the discharges around the mean firing interval [48].

### 4.4 Implementation remarks

The implementation of the main steps presented in the previous section can be summarized as follows. Once the matrices *S, A* and *B* are computed, the matrix *K* is determined using Eq. (6) by solving *n*_*e*_ linear systems. Then, Eq. (7) is used to find the solutions for *n*_*s*_ basis points, which is a fast matrix multiplication operation. For any given point source location ***r***, we compute its barycentric coordinates in associated tetrahedron and apply Eq. (5) to get values of electrical potentials at electrode locations. Finally, for a given fiber geometry, the single fiber action potential as recorded by the EMG electrodes is computed using Eq. (9).

The results presented in this study are obtained using a Python implementation of the proposed strategy. Assembling the matrix *A* and solving the system (6) is delegated to the FEniCS computing platform [49, 50]. The forearm geometry that is here representatively used as a conductor model is taken from the website of BodyParts3D, The Database Center for Life Science (http://lifesciencedb.jp/bp3d/). The volume mesh is generated from the surface meshes of the forearm tissues using the CGAL C++ library [51].

### 4.5 Computational performance

In this section, we report the computational time of the proposed model for a specific simulation case. The exact computational time values strongly depend on the implementation, experiment design, model parameters etc. The order of magnitude, however, stays the same. Note, that no multiprocessing tools were used in these computations. Each step, however, is highly scalable and can be efficiently distributed between parallel processes, which would significantly increase the performance. Computations for each muscle and fiber are independent and can be performed in parallel. Parallel computing would also apply to the electrodes in the general basis points computation.

For the purpose of demonstration, we simulated a 1-min-long, 100% maximum voluntary contraction (MVC) excitation of the Brachioradialis muscle with 50000 individual fibers and 200 motor units. The mesh of the volume conductor contained 2.1M vertices, which formed 13M tetrahedra. 16 rectangular and 16 circular electrodes were included in the model. The sampling frequency of the simulated signals was 2000 Hz. Table 1 shows the computational time for each of the main steps in this simulation.

**Table 1:**
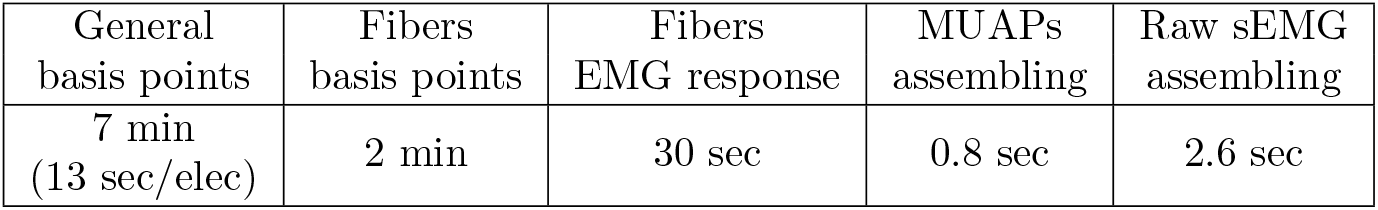
Computational performance of each of the main steps of a raw EMG simulation. General basis points computation refers to equation (7); fiber basis points are computed with equation (5); fibers EMG response is computed with equation (9).

An important property of our model is that each step depends only on the data produced by the previous steps. This property can be exploited to change some simulation parameters without recomputing every step of the simulation. For example, it is not necessary to recompute solutions for the fiber basis points if fibers geometry and conductor model stay the same and only the parameters related to the fiber properties (AP velocity, end-plate location, tendon sizes, etc.), MU distribution or recruitment model are modified. In this example, the **total** simulation time for this new set of parameters will only take approximately 30 + 0.8 + 2.6 = 33.4 s.

A brief description of the main parameters required at each step follows. The full arm and electrode geometry as well as the tissue conductivities define the computation of general basis points. To compute fibers basis points solutions, the 3D geometry of the fibers is required. Computing the fiber EMG responses requires the shape of the intracellular AP waveforms, AP propagation velocity, sizes of tendon and active fiber parts, neuromuscular junction location, fiber diameter and intracellular conductivity, and sampling frequency. To compute the MUs action potentials, the MU distribution in the muscle, i.e. the association of fibers to each motor unit, need to be defined. In the proposed model, once the number of MUs, their sizes and territory areas are selected, the MU distribution is randomly generated. Finally, to synthesize the sEMG signal, the muscle excitation drives and recruitment model parameters (motor unit recruitment thresholds and firing rates) are required.

### 4.6 Comparison with the cylindrical analytical solution

First, we compared our numerical solution with its analytical counterpart for a simple volume conductor geometry [21]. We used a four layer cylindrical model with layers corresponding to bone (*r* = 0.7cm), muscle (*r* = 2cm), fat(*r* = 2.3cm) and skin (*r* = 2.4cm) surfaces. 16 point electrodes were simulated on the skin surface directly above a fiber. The fiber was located at varying depths into the muscle tissue, in the range 1 mm to 11 mm. Differential sEMG signals were simulated using the analytical and numerical solutions of the forward problem.

### 4.7 Details of realistic simulation examples

For the single muscle excitation example, 50k muscle fibers were generated inside the muscle and distributed within 200 motor units. The size of MUs varied exponentially from 11 to 1150 fibers. The areas of MU territories varied from 10% to 50% of the muscle cross-sectional area. The muscle excitation drive was decomposed into MU impulse trains according to the size principle. In this example, the firing rate for each MU ranged from 8 Hz to 35 Hz and all MUs were recruited when an excitation level of 75% MVC was reached.

For the multiple muscles experiment, the flexor group included the Palmaris longus, Flexor carpi ulnaris (ulnar head), Flexor carpi ulnaris (humeral head), and Flexor carpi radialis muscles. The extensor group included the Extensor digitorum, Extensor carpi ulnaris, Extensor carpi radialis brevis, and Extensor carpi radialis longus muscles. During a wrist flexion, the muscles of the flexor group reached an excitation level of 90% MVC. During extension, extensor group was activated with the same exciation level. Moreover, a small but constant excitation of the abduction muscle group was added to simulate the wrist resistance against gravity. The abduction muscle group included the Flexor carpi radialis, Extensor carpi radialis brevis, and Extensor carpi radialis longus muscles. For each muscle, a number of muscle fibers between 32k and 78k was simulated, depending on the muscle cross-sectional area. Muscle fibers were distributed within motor units, whose number varied from 150 to 300 per muscle.

### 4.8 Details of deep learning experiment

To evaluate the effect of using the simulation-pre-trained network, an experimentally collected high-density surface electromyography (HD-sEMG) signal dataset was used, originally created to test wrist-wearable interfaces [52]. The experimental protocol was designed in agreement with the Declaration of Helsinki and was approved by Imperial College London ethics committee (JRCO: 18IC4685). Nine participants (4 females, 5 males, ages: 23-31) took part in the study after signing informed consent forms. The participants performed 5-second isometric contractions of their dominant-hand index finger at 15% of maximal force, with sEMG activity measured using two flexible 5×13 electrode grids with 8-mm spacing placed on the circumference of the wrist, immediately proximal to the ulnar head. HD-sEMG signal was sampled at 2048Hz, whilst force profiles were sampled with a custom load cell at 10Hz. The signal was then decomposed into motor neuron activity using convolutive blind source separation [53]. For the purpose of training and testing the supervised decomposition pipeline, motor neuron activity was accepted if it was present for at least 80% of the contraction window. For each participant the HD-sEMG signal and accompanying decomposed motor neuron activity (as a sparse binary matrix) was then split into a 4 second training window and a 1 second testing window.

A gated recurrent unit (GRU) network was used as the deep learning model due to previous studies showing good performance with this data type [26]. After hyperparameter optimisation by grid search, a minimally-parameterised model was found to perform optimally, likely due to the short length of the training data available. Input HD-sEMG signal was first encoded by a single layer GRU with a hidden dimension of 1024 in length [54]. To make a time instant prediction a densely-connected linear layer with sigmoid activation function took as an input a moving 20 sample-wide window from the GRU output, centred on the time instant of interest. Predicted activity was converted to spike timestamps using a two-class K-means clustering algorithm. Binary cross entropy was used as the loss function and Adam with weight decay used as the optimising algorithm [55].

To improve model generalisation an early-stopping framework was used, based on 10% of the training data retained as a validation set. Training, validation and test data was z-score standardised using the mean and standard deviation calculated from the training set. During training the input signal was augmented with noise of standard normal distribution. To account for the high sparsity of the output matrix, samples containing motor neurons were artificially oversampled, with each each input batch of 512 time instants containing at least 20% motor neuron activation. All machine learning was implemented using the pytorch library in python. Final performance was assessed using the rate of agreement metric (RoA).

The optimised architecture of the GRU network was used for pre-training, which was conducted using multi-task learning in a hard parameter sharing paradigm [56]. Four digital twins were created for simulation using different model parameters (tissue conductivities, MU distribution, fiber properties, etc.), with the generated motor unit activation (MUAP) templates from flexor digitorum profundus and superficialis used to create 64 sets, each containing 5 MUAPs. Each set was used to generate windows of signal with a range of MUAP superpositions (Fig. 6A). In signal windows with motor neuron activity a MUAP template was placed in the centre of the window, before being additively superimposed with a random number of MUAP templates from other motor units at random time offsets. In windows without activity no template was placed in the centre of the window. During multi-task learning training, the same GRU layer (and parameters) were shared between the 64 recordings, but each recording had its own output layer, operating on a 20 sample-wide window as in the experimental recordings (Fig. 6B). In this way the GRU layer was trained to act as a more general feature extractor, whilst the individual linear output layers made class predictions specific to each recording. Training again used noise augmentation, binary cross-entropy and Adam with weight decay.

**Figure 6:**
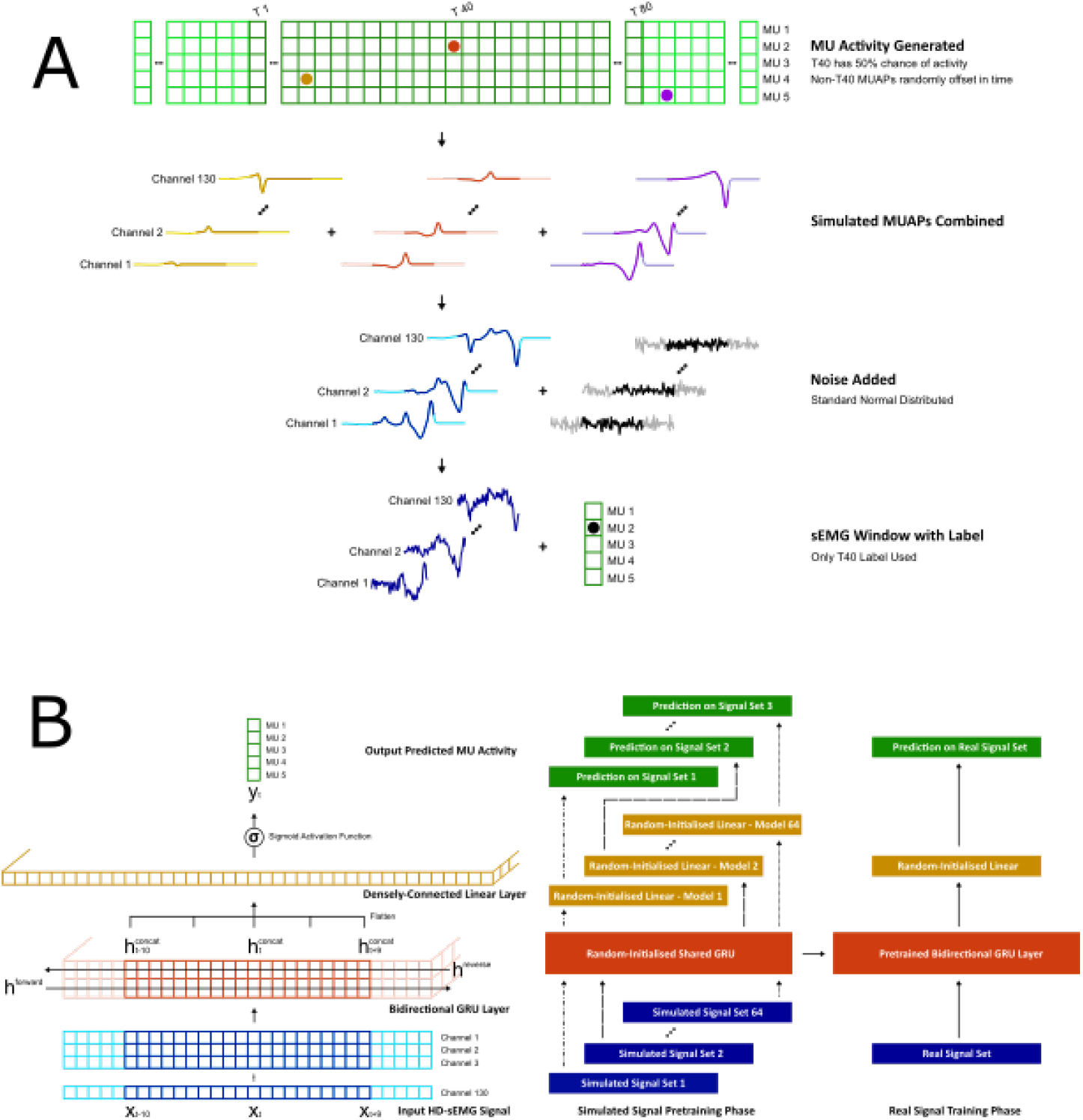
(A) Methodology used to build windows from the simulated MUAP template set for the pre-training phase. Each simulated template was 160 samples wide at a 2048Hz sampling rate and with 130 channels. First either a MUAP template was placed in the centre of the window or it was left empty at a 50% probability. Then MUAP templates from other MU classes were added to the window at a random offset to generate superpositions. Finally standard normal distributed noise was added to the window, with the central 80 samples then paired with the label for supervised learning. (B) The neural network architecture and pre-training methodology used to improve the performance of a deep learning-based HD-sEMG decomposition algorithm. The neural network consists of a single gated recurrent unit layer, with predictions made using a 20-sample wide window of the hidden vector output, which is flattened before being passed to a sigmoid-activated densely-connected linear layer. In the pretraining phase a multi-task learning regimen is used to optimise the parameters of the gated recurrent unit using the simulated sEMG. This pre-trained layer can then be used to improve the optimisation performance on real sEMG data.

To use the simulation-pre-trained network in the experimental data the GRU parameters from the pre-trained network were used, whilst the linear output layer used a normal random initialisation. This was the compared to a normal random initialisation of both the GRU and output layer. In both instances the network was trained using the methodology specified above, with the only difference being whether the GRU layer was simulation-pre-trained or not.

## Acknowledgments

For this study, DF was sponsored by the European Research Council (ERC) under the Synergy Grant Natural BionicS (810346) and the EPSRC Transformative Healthcare for 2050 project NISNEM Technology (EP/T020970/1). AC and IMG are sponsored by the Engineering and Physical Sciences Research Council (EPSRC) - Centre for Doctoral Training in Neurotechnology for Life and Health and Meta.

## Author contributions

KM, SDG, and DF conceptualized the study. KM and SDG developed the software implementation of the simulator. KM, AC, IMG, SDG, and DF performed the experimental measures and conceptualized the data analysis. AC and IMG performed the data analysis. KM, SDG, and DF prepared the first draft of the manuscript. All authors edited the manuscript for important scientific content and all approved the final version.

## References

[1] Farina, D. et al. Toward higher-performance bionic limbs for wider clinical use. Nature Biomedical Engineering (2021).

[2] Farina, D. et al. The extraction of neural information from the surface EMG for the control of upper-limb prostheses: Emerging avenues and challenges. IEEE Transactions on Neural Systems and Rehabilitation Engineering 22, 797–809 (2014).

[3] Farina, D. et al. Man/machine interface based on the discharge timings of spinal motor neurons after targeted muscle reinnervation. Nature Biomedical Engineering 1, 0025 (2017).

[4] Zhuang, K. Z. et al. Shared human–robot proportional control of a dexterous myoelectric prosthesis. Nature Machine Intelligence 1, 400–411 (2019).

[5] Geng, W. et al. Gesture recognition by instantaneous surface EMG images. Scientific Reports 6, 36571 (2016).

[6] Guo, W. et al. Long exposure convolutional memory network for accurate estimation of finger kinematics from surface electromyographic signals. Journal of Neural Engineering 18, 026027 (2021).

[7] Guerra, I. M., Barsakcioglu, D. Y., Vujaklija, I., Wetmore, D. Z. & Farina, D. Far-field electric potentials provide access to the output from the spinal cord from wrist-mounted sensors. Journal of Neural Engineering (in press). URL https://www.biorxiv.org/content/10.1101/2021.04.06.438640v1.

[8] Bird, J. J., Pritchard, M., Fratini, A., Ekart, A. & Faria, D. R. Synthetic biological signals machine-generated by GPT-2 improve the classification of EEG and EMG through data augmentation. IEEE Robotics and Automation Letters 6, 3498–3504 (2021).

[9] Tsinganos, P., Cornelis, B., Cornelis, J., Jansen, B. & Skodras, A. Data augmentation of surface electromyography for hand gesture recognition. Sensors (Switzerland) 20, 4892 (2020).

[10] Wang, F., Zhong, S.-h., Peng, J., Jiang, J. & Liu, Y. Data augmentation for EEG-based emotion recognition with deep convolutional neural networks.In Schoeffmann, K. et al. (eds.) MultiMedia Modeling, 82–93 (Springer International Publishing, Cham, 2018).

[11] Zanini, R. A. & Colombini, E. L. Parkinson’s disease EMG data augmentation and simulation with DCGANs and style transfer. Sensors (Switzerland) 20, 2605 (2020).

[12] Wen, S. et al. Rapid adaptation of brain–computer interfaces to new neuronal ensembles or participants via generative modelling. Nature Biomedical Engineering (2021).

[13] Gootzen, T. H. J. M., Stegeman, D. F. & van Oosterom, A. Finite limb dimensions and finite muscle length in a model for the generation of electromyographic signals. Electroencephalography and Clinical Neurophysiology/ Evoked Potentials 81, 152–162 (1991).

[14] Fuglevand, A. J., Winter, D. A., Patla, A. E. & Stashuk, D. Detection of motor unit action potentials with surface electrodes: influence of electrode size and spacing. Biological Cybernetics 67, 143–153 (1992).

[15] Stegeman, D. F. & Linssen, W. H. Muscle fiber action potential changes and surface EMG: A simulation study. Journal of Electromyography and Kinesiology 2, 130–140 (1992).

[16] Yue, G., Fuglevand, A. J., Nordstrom, M. A. & Enoka, R. M. Limitations of the surface electromyography technique for estimating motor unit synchronization. Biological Cybernetics 73, 223–233 (1995).

[17] Roeleveld, K., Blok, J. H., Stegeman, D. F. & Oosterom, A. V. Volume conduction models for surface emg; confrontation with measurements. Journal of Electromyography and Kinesiology 7, 221–232 (1997).

[18] Schneider, J., Silny, J. & Rau, G. Influence of tissue inhomogeneities on noninvasive muscle fiber conduction velocity measurements—investigated by physical and numerical modeling. IEEE Transactions on Biomedical Engineering 38, 851 – 860 (1991).

[19] Botelho, D. P., Curran, K. & Lowery, M. M. Anatomically accurate model of EMG during index finger flexion and abduction derived from diffusion tensor imaging. PLoS Computational Biology 15, 1–24 (2019).

[20] Vecchio, A. D. D. et al. Spinal motoneurons of the human newborn are highly synchronized during leg movements. Science Advances 6, eabc3916 (2020).

[21] Farina, D., Mesin, L., Martina, S. & Merletti, R. A surface EMG generation model with multilayer cylindrical description of the volume conductor. IEEE Transactions on Biomedical Engineering 51, 415–426 (2004).

[22] Merletti, R. & Muceli, S. Tutorial. Surface EMG detection in space and time: Best practices. Journal of Electromyography and Kinesiology 49, 102363 (2019).

[23] Henneman, E. Relation between size of neurons and their susceptibility to discharge. Science 126, 1345–1347 (1957).

[24] Carriou, V., Boudaoud, S., Laforet, J. & Ayachi, F. S. Fast generation model of high density surface EMG signals in a cylindrical conductor volume. Computers in Biology and Medicine 74, 54–68 (2016).

[25] Girshick, R., Donahue, J., Darrell, T. & Malik, J. Rich feature hierarchies for accurate object detection and semantic segmentation. In Proceedings of the IEEE conference on computer vision and pattern recognition, 580–587 (2014).

[26] Clarke, A. K. et al. Deep learning for robust decomposition of high-density surface EMG signals. IEEE Transactions on Biomedical Engineering 68, 526 – 534 (2021).

[27] Merletti, R. & Farina, D. Surface Electromyography : Physiology, Engineering, and Applications (John Wiley & Sons, Ltd, 2016).

[28] Dimitrov, G. V. & Dimitrova, N. A. Precise and fast calculation of the motor unit potentials detected by a point and rectangular plate electrode. Medical Engineering and Physics 20, 374–381 (1998).

[29] Farina, D. & Merletti, R. A novel approach for precise simulation of the EMG signal detected by surface electrodes. IEEE Transactions on Biomedical Engineering 48, 637–646 (2001).

[30] Plonsey, R. Action potential sources and their volume conductor fields. Proceedings of the IEEE 65, 601–611 (1977).

[31] Plonsey, R. & Heppner, D. B. Considerations of quasi-stationarity in electrophysiological systems. The Bulletin of mathematical biophysics 29, 657—664 (1967).

[32] Heringa, A., Stegeman, D. F., Uijen, G. J. & Weerd, J. P. D. Solution methods of electrical field problems in physiology. IEEE Transactions on Biomedical Engineering BME-29, 34–42 (1982).

[33] Farina, D., Mesin, L. & Martina, S. Advances in surface electromyographic signal simulation with analytical and numerical descriptions of the volume conductor. Medical and Biological Engineering and Computing 42, 467 (2004).

[34] Nicholson, C. & A. Freeman J. Theory of current source density analysis and determination of conductivity tensor for anuran cerebellum. Journal of Neurophysiology 38, 356–368 (1975).

[35] Pettersen, K. H., Lindén, H., Dale, A. M. & Einevoll, G. T. Extracellular spikes and current-source density, 92–135 (Cambridge University Press, Cambridge, UK, 2010).

[36] Peter Knabner, L. A. The Finite Element Method for the Poisson Equation, 46–91 (Springer New York, New York, NY, 2003).

[37] Vallaghé, S., Papadopoulo, T. & Clerc, M. The adjoint method for general EEG and MEG sensor-based lead field equations. Physics in Medicine and Biology 54, 135–147 (2008).

[38] Plonsey, R. The active fiber in a volume conductor. IEEE Transactions on Biomedical Engineering BME-21, 371 – 381 (1974).

[39] Rosenfalck, P. Intra- and extracellular potential fields of active nerve and muscle fibres. A physico-mathematical analysis of different models. Acta physiologica Scandinavica. Supplementum 321, 1—168 (1969).

[40] Burke, R. E., Levine, D. N., Tsairis, P. & Zajac, F. E. Physiological types and histochemical profiles in motor units of the cat gastrocnemius. The Journal of Physiology 234, 723–748 (1973).

[41] Schiaffino, S. & Reggiani, C. Fiber types in mammalian skeletal muscles. Physiological Reviews 91, 1447–1531 (2011).

[42] Modenese, L. & Kohout, J. Automated generation of three-dimensional complex muscle geometries for use in personalised musculoskeletal models. Annals of Biomedical Engineering 48, 1793–1804 (2020).

[43] Keenan, K. G., Farina, D., Merletti, R. & Enoka, R. M. Influence of motor unit properties on the size of the simulated evoked surface EMG potential. Experimental Brain Research 169, 37–49 (2006).

[44] Carriou, V., Laforet, J., Boudaoud, S. & Al Harrach, M. Realistic motor unit placement in a cylindrical HD-sEMG generation model. In 2016 38th Annual International Conference of the IEEE Engineering in Medicine and Biology Society (EMBC), 1704–1707 (IEEE, Orlando, United States, 2016). URL https://hal.archives-ouvertes.fr/hal-03586013.

[45] Fuglevand, A., Winter, D. A. & Patla, A. E. Models of recruitment and rate coding organization in motor-unit pools. Journal of Neurophysiology 70, 2470–2488 (1993).

[46] Ayachi, F. S., Boudaoud, S. & Marque, C. K. Evaluation of muscle force classification using shape analysis of the sEMG probability density function: A simulation study. Medical and Biological Engineering and Computing 52, 673–684 (2014).

[47] Luca, C. J. D. & Hostage, E. C. Relationship between firing rate and recruitment threshold of motoneurons in voluntary isometric contractions. Journal of Neurophysiology 104, 1034–1046 (2010).

[48] Arabadzhiev, T. I., Dimitrov, V. G., Dimitrova, N. A. & Dimitrov, G. V. Influence of motor unit synchronization on amplitude characteristics of surface and intramuscularly recorded EMG signals. European Journal of Applied Physiology 108, 227 (2010).

[49] Logg, A., Mardal, K. A. & Wells, G. N. Automated solution of differential equations by the finite element method, vol. 84 LNCSE of Lecture Notes in Computational Science and Engineering (Springer, Berlin, Heidelberg, 2012).

[50] Alnæs, M. et al. The FEniCS Project Version 1.5. Archive of Numerical Software 3 (2015).

[51] The CGAL Project. CGAL User and Reference Manual (CGAL Editorial Board, 2021), 5.2.1 edn. URL https://doc.cgal.org/5.2.1/Manual/packages.html.

[52] Guerra, I. M., Barsakcioglu, D. Y., Vujaklija, I., Wetmore, D. Z. & Farina, D. Non-invasive real-time access to the output of the spinal cord via a wrist wearable interface. bioRxiv (2021).

[53] Negro, F., Muceli, S., Castronovo, A. M., Holobar, A. & Farina, D. Multichannel intramuscular and surface EMG decomposition by convolutive blind source separation. Journal of Neural Engineering 13, 026027 (2016).

[54] Cho, K. et al. Learning phrase representations using RNN encoder-decoder for statistical machine translation. arXiv preprint arXiv:1406.1078 (2014).

[55] Loshchilov, I. & Hutter, F. Decoupled weight decay regularization. arXiv preprint arXiv:1711.05101 (2017).

[56] Baxter, J. A Bayesian information theoretic model of learning to learn via multiple task sampling. Machine learning 28, 7–39 (1997).

